# Deep Transfer Learning of Drug Responses by Integrating Bulk and Single-cell RNA-seq data

**DOI:** 10.1101/2021.08.01.454654

**Authors:** Junyi Chen, Zhenyu Wu, Ren Qi, Anjun Ma, Jing Zhao, Dong Xu, Lang Li, Qin Ma

**Affiliations:** Department of Biomedical Informatics, College of Medicine, The Ohio State University, Columbus, OH 43210, USA; Department of Electrical Engineering and Computer Science, and Christopher S. Bond Life Sciences Center, University of Missouri, Columbia, MO 65211, USA

**Keywords:** Deep transfer learning, drug response prediction, single-cell RNA-seq, integrative analysis

## Abstract

Massively bulk RNA sequencing databases incorporating drug screening have opened up an avenue to inform the optimal clinical application of cancer drugs. Meanwhile, the growing single-cell RNA sequencing (scRNA-seq) data contributes to improving therapeutic effectiveness by studying the heterogeneity of drug responses for cancer cell subpopulations. There is a clear significance in developing computational biology approaches to predict and interpret cancer drug response in single cell data from clinical samples. Here, we introduce scDEAL, a deep transfer learning framework for cancer drug response prediction at single-cell level by integrating large-scale bulk cell line data. The true innovation of scDEAL is to translate cancer cell line drug responses into predicting clinical drug responses via learning relations of gene expressions and drug responses at bulk-level and transfer to predict drug responses in scRNA-seq. Another innovation is the integrated gradient feature interpretation to infer a comprehensive set of signature genes to reveal potential drug resistance mechanisms. We benchmarked scDEAL on six scRNA-seq datasets and indicate its model interpretability through these case studies. We believe that this work may help study cell reprogramming, drug selection, and repurposing for improving therapeutic efficacy.

## Background

The investigation and development of precision medicine have achieved remarkable progress in understanding the complexity of the genomic landscape of cancer. The expectation towards tailoring cancer treatments to a particular genomic signature of an individual cell is growing rapidly. Abundant *in vitro* drug screening studies have been conducted, giving rise to drug response data on different cancer cell lines (CCLs) (1,2). Yet, one obstacle in cancer drug treatments is the low efficacies and high relapse rates caused by cancer heterogeneity among different states or cell fates. Such heterogeneity is responsible for differentiated responses of individual cells to a drug, leading to minimal residues remaining in the body, and finally, cancer relapse (3). Single-cell RNA-sequencing (scRNA-seq) technique provides an unprecedented opportunity to discover heterogeneous gene expressions of cancer subpopulations in response to specific drugs (4). All existing drug response prediction methods were developed for bulk data that cannot be directly used for larger-scale and highly intricate single-cell data. Hence, it is very much needed to develop computational methods to infer heterogeneous drug responses at the single-cell level.

Deep learning methods have been deployed to tackle scRNA-seq data, achieving ideal performances in abstracting low-dimensional features and recovering dropout issues (5,6). The main obstacle in developing a deep learning-based tool for predicting single-cell drug responses is the lack of sufficient training power due to the limited number of benchmarked data in the public domain. As far as we know, only six datasets with drug treatment and experimentally validated drug responses for individual cells are available in the public domain. Fortunately, bulk drug-related RNA-seq data can be great complementary resources to infer relations of gene expression-drug response to help predict drug responses at the single-cell level (7), if bulk and single-cell data can be integrated (8).

Deep transfer learning (DTL) is a deep learning method with the aim to transfer knowledge from one model to another (9). Using DTL, we can solve a particular task using a whole or part of an already pre-trained model on a different task. It has been applied as an effective strategy in leveraging multiple bulk data sources for cancer drug response predictions (10), yet, its power in transferring valuable bulk-level knowledge to the single-cell level has not been investigated. To this end, we developed a DTL framework, named scDEAL (<u>s</u>ingle-<u>c</u>ell <u>D</u>rug r<u>E</u>sponse <u>A</u>na<u>L</u>ysis), to predict drug responses of specific cancer cell groups from scRNA-seq data. To the best of our knowledge, scDEAL is the first computational framework that integrates a broad range of bulk and scRNA-seq data for drug response prediction. scDEAL holds a strong prediction power in predicting single-cell level drug sensitivity since it establishes bridges among drug sensitivity, gene features in single cells, and gene features in bulk samples. Corresponding parameters were trained and optimized towards the most reliable way to transfer information via the bridges (11), which is pre-trained (12) on a large volume of bulk cell line data stored in the Genomics of Drug Sensitivity in Cancer (GDSC) database (13,14). Applied on six benchmark drug-treated scRNA-seq data, scDEAL achieves confidently high accuracy in predicting cell-type drug responses, comparing to benchmark labels. We further identified and interpreted critical genes that are either responsible for the development of sensitivity or resistance in a cell, by tracing and accumulating integrated gradients of each neuron in the neural network. Last, we demonstrated the relations between the dynamic change of drug response among cells and cell developmental trajectory, providing clues for identifying minimal residue disease cells.

## Results

### Overview of the scDEAL framework

In general, scDEAL models relations between the gene expression feature and drug response at the bulk level, where annotations for cell lines are accessible (**Fig. 1A**). It deducts the single-cell expression features, having the maximum association with expression features at the bulk level. Therefore, the *single-cell expression–drug response* relations can be obtained through the migration of *bulk expression-drug response* relations. With the above strategies, scDEAL infers drug responses for individual cells without the training needs at the single-cell level.

**Fig. 1.**
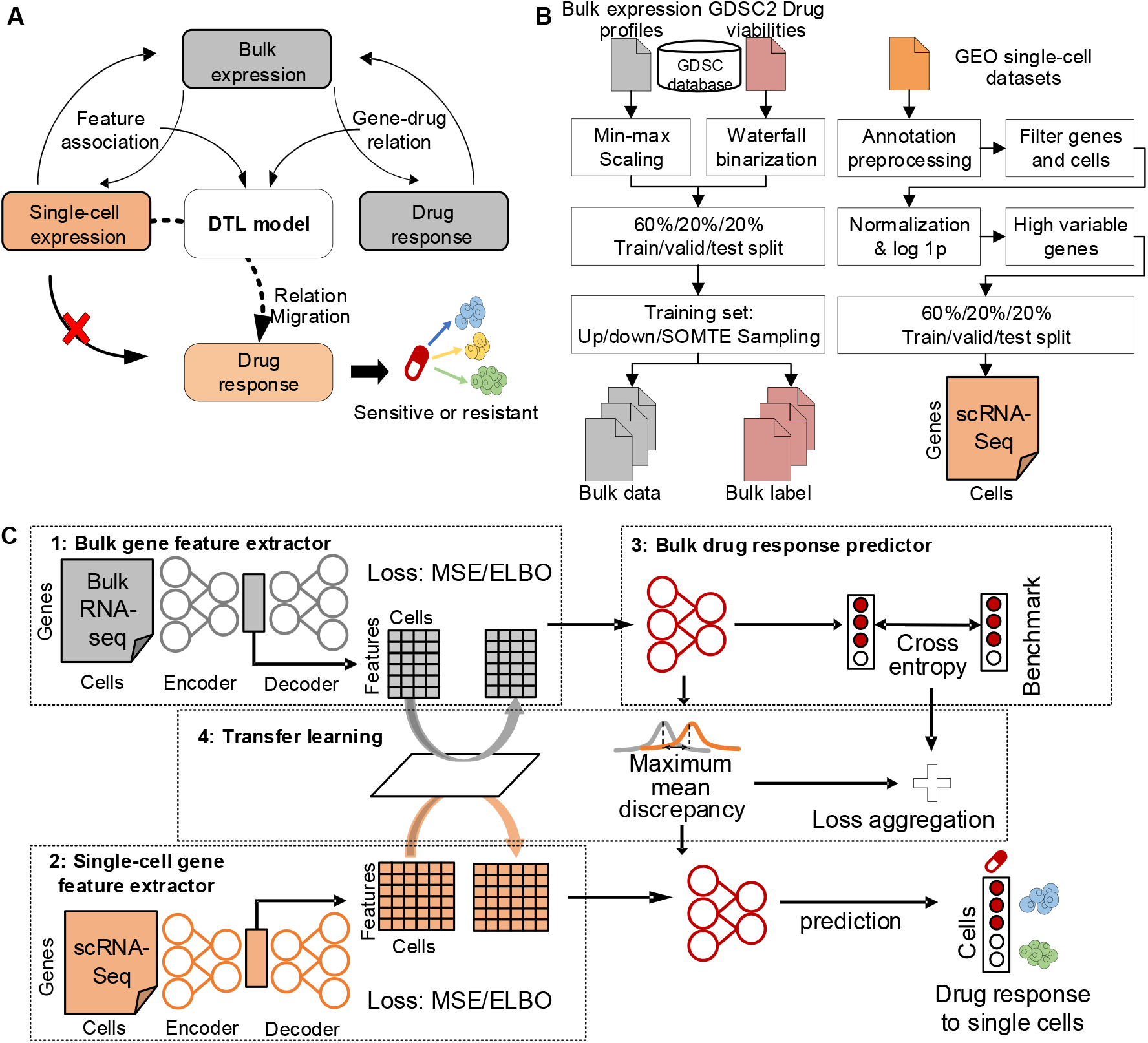
The workflow of scDEAL. (**A**) The strategy of scDEAL is to transfer the knowledge through the DTL model of two relations of (*i*) gene expression feature association between bulk RNA-seq and scRNA-seq, and (*ii*) genes and drug responses to predict drug responses in individual cell types. (**B**) Data pre-processing steps for GDSC bulk data and scRNA-seq data. (**C**) The scDEAL framework, with four components: 1) a bulk gene feature extractor, 2) a single-cell gene feature extractor, 3) a drug response predictor, and 4) a transfer learning model. Bulk and single-cell feature extractors extract genetic features from expression profiles. It is firstly trained in an encoder-decoder manner. Subsequently, the bulk gene features extracted by the drug response predictor are applied to train a fully connected predictor for drug response prediction. Finally, the transfer learning model is used to update two feature extractors and the predictor at once. It enables the predictor to received genes features extracted from scRNA-seq data to generate single-cell drug response prediction.

Both bulk and scRNA-seq data were pre-processed prior to the input of scDEAL (**Fig. 1B**). The scDEAL framework involves four major components: 1) the bulk gene feature extractor, 2) the single-cell gene feature extractor, 3) the drug response predictor, and 4) the whole DTL model combining all extractors and predictor as one (**Fig. 1C**). Two feature extractors (bulk and single-cell) are pre-trained in the encoder-decoder structure to extract low dimensional gene features from bulk and scRNA-seq data, respectively. The pre-training reduces the reconstruction loss between the decoder output and the expression profiles, making low dimensional features informative to represent the original gene expressions. Meanwhile, the pre-training is also a fine-tuned process (12) to generate initial better neuron weights within the DTL model. A fully connected predictor is attached to the pre-trained bulk feature extractor. It is trained to minimize the difference between the output and the drug response labels by the cross-entropy loss. The predictor attached to the extractor (extractor-predictor) is trained to receive bulk RNA-seq data and predict drug response labels. Ultimately, the DTL model updates feature extractors and the predictor simultaneously in a multi-task learning manner, receiving both bulk and scRNA-seq data as inputs. Specifically, the first task is to minimize the differences (mean maximum discrepancy loss) between gene features from two extractors, bridging the communication between bulk and scRNA-seq data. The second task is to re-minimize the cross-entropy loss of extractor-predictor output and the drug response, similar to the training of the predictor. The output of scDEAL is the prediction of the potential drug response of individual cells or cell clusters (if available).

### Training of scDEAL and assessment of prediction accuracy

The training of scDEAL is composed of a source model trained to predict bulk-level drug response and a target model trained with DTL to predict single-cell level response. We evaluated the DTL performance of scDEAL on a collection of six published datasets covering five drug treatments, including Cisplatin, Gefitinib, I-BET-762, Docetaxel, and Erlotinib. Here, we focus on the accuracy of single-cell level response predictions while bulk level benchmarks are presented **(Supplementary Table S1**). All selected datasets have been provided with drug response annotations (i.e., drug sensitive or resistant) for individual cells. The prediction of drug responses in scDEAL is compared to the ground truth using three commonly applied metrics for machine learning: F-score, Area Under the Receiver Operating Characteristic (**AUROC**), and average precision (**AP**). **Fig. 2A** presents the results of scDEAL measured by three metrics on six datasets. As a result, scDEAL achieves high prediction performance across different datasets and metrics. In particular, ROC and F-score are more significant than 0.9 in Kong’s dataset (15) (GSE112274), and Schnepp dataset (16) (GSE140440), as well as AP is grated than 0.8 in the Sharma (HN120) dataset (17) (GSE118782). The excellent matching of scDEAL comparing with the ground truth cell types is also intuitive.

**Fig. 2.**
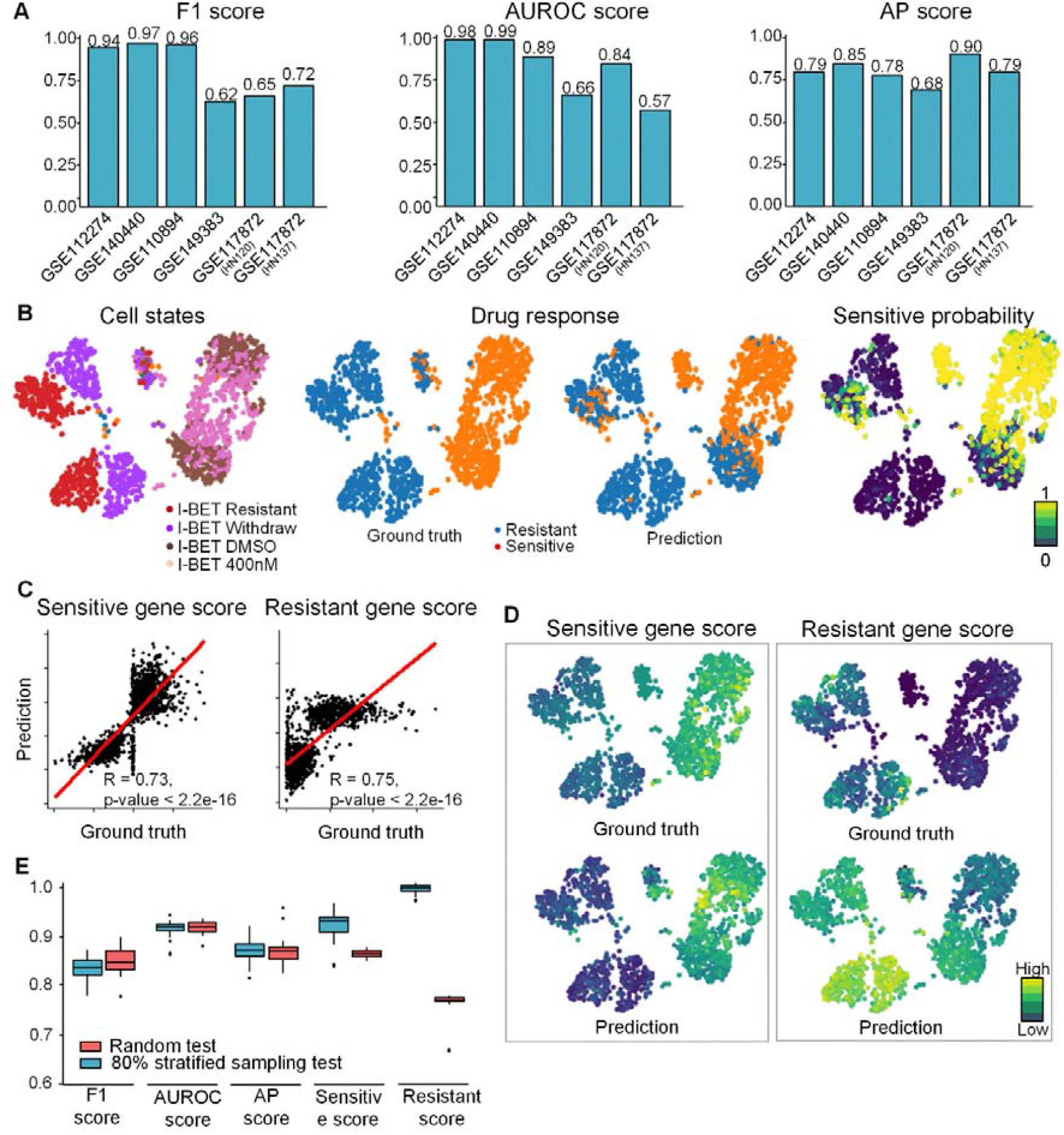
Benchmarking results of scDEAL. (**A**) Comparison of drug prediction performance across five datasets of scDEAL, measured by F-score, AUROC, and AP. (**B**) Cell embedding visualization based on UMAP of data from GSE110894. Different colors present the ground truth sensitivity, the prediction results, prediction probability, and cell types. (**C**) Linear regression plot displaying the relationship between the gene score derived from the predicted and the ground truth cell labels. (**D**) UMAP plot colored by sensitive (and resistant) gene score derived from differentially expressed genes in the predicted and ground truth sensitive (and resistant) cluster. (**E**) Box plot across five benchmark scores in random repetition test and 80% stratified sampling test.

To illustrate detailed predictions, we showcased the analytical power of scDEAL on Bell’s dataset (18) (GSE110894) containing 1,419 leukemic cells treated with BET inhibitor **(Fig. 2B)**. Four cell states including two sensitive states “dosed with DMSO” and “sensitive to 400 nM of IBET”, and two resistant states “cells have >IC90 to 1000 nM of IBET” and “withdrawn from IBET four days before experiments” (18). The prediction of scDEAL derived a consistent conclusion on leukemic cell drug responses with the original study. In all, 609 out of 685 resistant cells (>IC90 to 1000 nM of IBET) and the prolonged drug withdrawal cells are classified as resistant cells via scDEAL (**Fig. 2B**). Also, the sensitive prediction probability of cells is highly correlated to the ground truth label.

Though scDEAL delivered decent results on metrics comparing predictions and binarized drug response labels. The comparison cannot reflect the expression patterns of MA9 leukemic cells and my struggle in cells having sensitive probabilities in the borderline. To this end, we introduced a gene score reflecting drug responses (sensitivity or resistance) based on the differentially expressed genes in the sensitive (or resistant) cell cluster. The hypothesis behind the score is that an accurate prediction assigns the correct response label to cells. Therefore, the marker gene sets between the resistant and the sensitive state for an accurate prediction should be similar to the set derived from the ground truth. Gene scores based on the predicted drug response are highly correlated to that ground truth (**Fig. 2C**). The prediction and benchmark of both sensitive and resistance scores showed similar distributions on the UMAP, also indicating the performance if scDEAL **(Fig. 2D**).

Random repetition and stratified sampling were used to test the reproducibility of scDEAL (**Fig. 2E**). As a result, the variation of F-score, AUPRC, AP score, sensitivity score, and resistance score are 0.022, 0.017, and 0.022, respectively (**Supplementary Table S2**), indicating that scDEAL is robust across multiple runs of random sampling.

### Predicting critical genes for drug responsiveness using integrated gradient

Though scDEAL delivered accurate prediction for single-cell drug responses with the help of the DTL, comprehension of active genetic features within the model is essential to be revealed. To this end, we introduced Integrated Gradients (**IG**), a state-of-the-art feature interpretation method for deep neural networks, to provide a scope of “critical genes” contributing to the prediction of drug responses. Critical genes are defined based on the higher accumulation of gradients of neurons in the neural network following the path of layer connections. We conducted the IG analysis on the scDEAL model for the oral squamous cell carcinoma (**OSCC**) treated by Cisplatin in Sharma’s dataset (GSE118782) (17), characterizing the relations between transcriptional expressions and resistant/sensitive states. Using scDEAL, 85% of cells were correctly predicted as either sensitive or resistant cells to Cisplatin, with an F1-score of 0.74, AUROC score of 0.89, and AP score of 0.94 (**Fig. 3A**). The Pearson’s correlations of sensitive and resistant cells between prediction and benchmark are 0.83 and 0.82, respectively. We identified 714 sensitive critical genes and 364 resistant critical genes, according to the distinct patterns in the predicted clusters based on the gradients (**Fig. 3B**), which cannot be reflected directly from gene expressions (**Supplementary Fig. S1**).

**Fig. 3.**
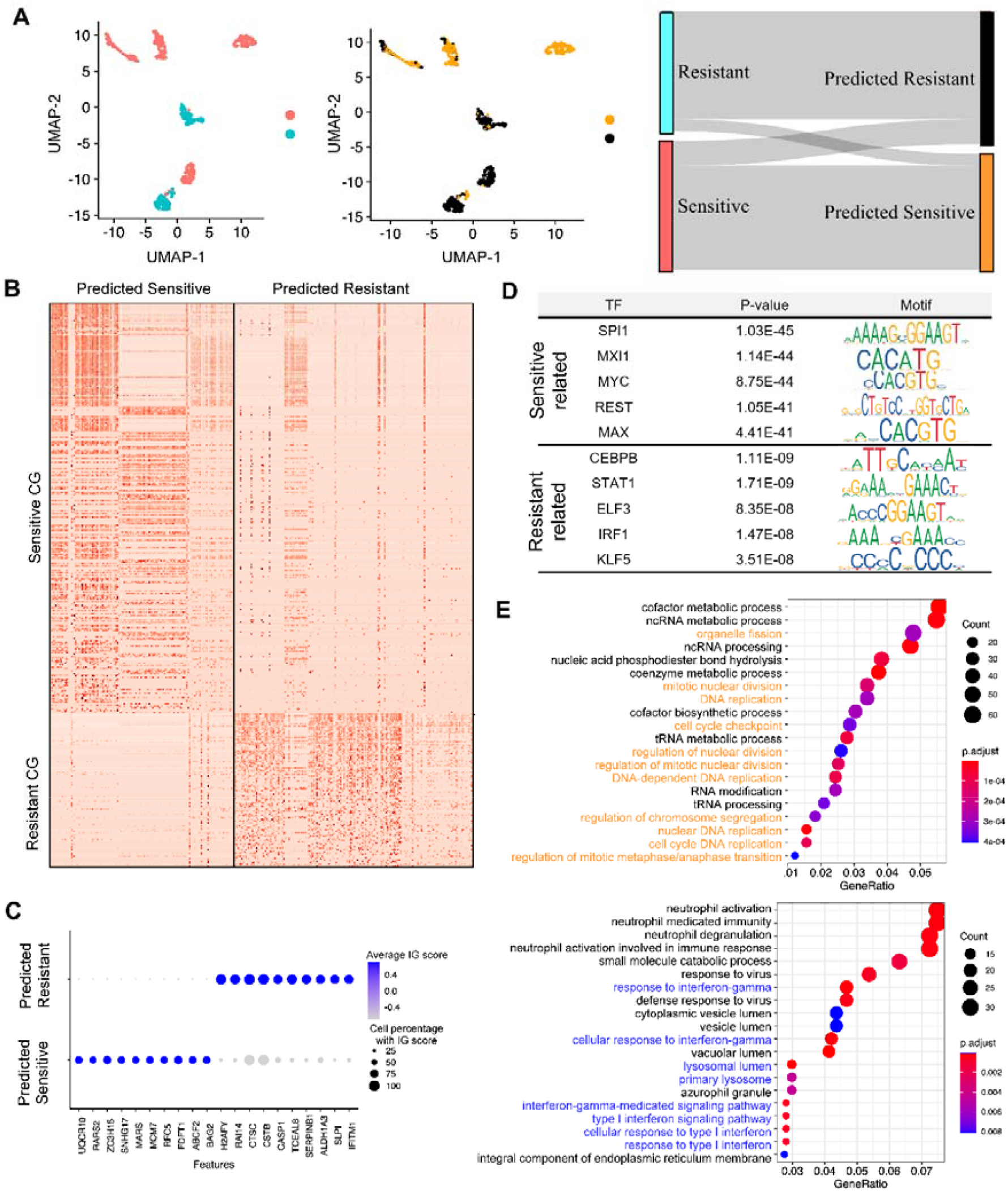
Case study of scDEAL using the Sharma (HN120) dataset **(A)** Response predictions of the dataset. The red and blue labels are ground truth, while the black and orange labels are from prediction. **(B)** The integrated gradient heatmap of critical genes in the predicted sensitive and resistant clusters. **(C)** The integrated gradient of identified critical genes of crucial pathways. **(D)** The binding motif logos of TFs that regulate the critical genes. **(E)** The Gene Ontology analysis of sensitive (upper panel) and resistant (lower panel) critical genes. The orange and blue highlights are the known pathways related to the Cisplatin treatment.

Cisplatin is one of the most widely used drugs for treating solid cancers such as testicular, ovarian, head and neck, bladder, lung, cervical cancer, melanoma, lymphomas (19). Cisplatin exerts its anti-cancer activity via the generation of DNA lesions by interacting with purine bases on DNA, interfering with DNA repair and causing DNA damage, followed by activation of several signal transduction pathways and finally leading to apoptosis of cancer cells (20). Particularly, sensitive CGs MCM7 and RFC5 have already been reported to play a role in Cisplatin-related pathways, such as cell cycle control, DNA replication, and error repair (21,22). The down-regulation of MCM7 was reported as an indicator of the increase of Cisplatin resistance in bladder cancer (23). Resistant CGs CTSC, CSTB, and IFITM1 are key players in the IFN signal transduction (lysosome dependent apoptosis and E-cadherin signal) (24-26). We further checked the top 10 sensitive and resistant CGs and found no explicit expression patterns but distinct integrated gradients differences (**Fig. 3C** and **Supplementary Fig. S1**). We utilized LISA2 (27) to identify potential regulators of sensitive and resistant CGs (**Fig. 3D**). SPI1, MXI1, MYC, REST, and MAX are the potential regulators of sensitive CGs that are well known for their roles in cell cycle regulation. The down-regulation of MYC and cell cycle regulation is reported to increase Cisplatin resistance(28). On the other hand, five TFs were also predicted for resistant CGs, including CEBPB, STAT1, ELF3, IRF1, and KLF5. STAT1 and IRF1 in the potential regulators of resistant CGs indicate the participation of JAK-STAT signals, which are tightly working with the IFN signals. The IFN signal, along with the JAK-STAT signals, may promote Cisplatin resistance in OSCC.

We performed the Gene Ontology analysis for the sensitive and resistant critical genes, respectively. The sensitive CGs are primarily enriched in cell cycle and DNA replication-related pathways (**Fig. 3E**). It has also been reported that the mitochondria can be attacked by Cisplatin (29). ROS production (reactive oxygen species) triggered by the attack may explain the enriched carbon metabolism pathways in the KEGG analysis results. Thus, our sensitive CG indicates and well presents the potential working mechanisms of Cisplatin. On the other hand, increased expression levels of various transporters and increased repair of platinum-DNA adducts are considered the most significant processes in developing drug resistance (29). In addition, the autophagy, mitochondria quality control may establish the resistance, lysosomal degradation of organelle, and the defects in the signal transduction pathways that usually elicit apoptosis in response to DNA damage and problems with the cell death executioner machinery (30,31). Those resistance-related pathways are also shown in our KEGG and GO analysis of resistant CGs (**Fig. 3E** and **Supplementary Fig. S1**). Interestingly, we found the interferon (IFN) related pathway is highly enriched in our resistant CGs, as IFN family genes are not among the typical genes related to drug resistance. Studies have reported that the IFN-gamma may suppress the caspase-3 activation and the apoptosis induced by Cisplatin in renal cells (32). Overall, our KEGG and GO results are in accordance with the Cisplatin anti-tumor mechanisms mentioned above.

### Correlating drug responsiveness with pseudotime analysis

Genetic features of developing the drug resistance are characterized by the integrated gradient method on the trained scDEAL model. Using the same OSCC data, we further inspected scDEAL predictions and the characterized critical gene signals accompanied by the transcriptome dynamics and drug resistance developments. We hypothesized that the drug response in OSCC decreased along with cell states starting from sensitive to resistant. We applied PAGA (33) and **diffusion pseudotime** (34) as the trajectory inference to the OSCC data, covering four stages divided into six cell clusters (**Fig. 4A**) of tumor evolution under the selection pressure of Cisplatin and metastatic dissemination.

**Fig. 4.**
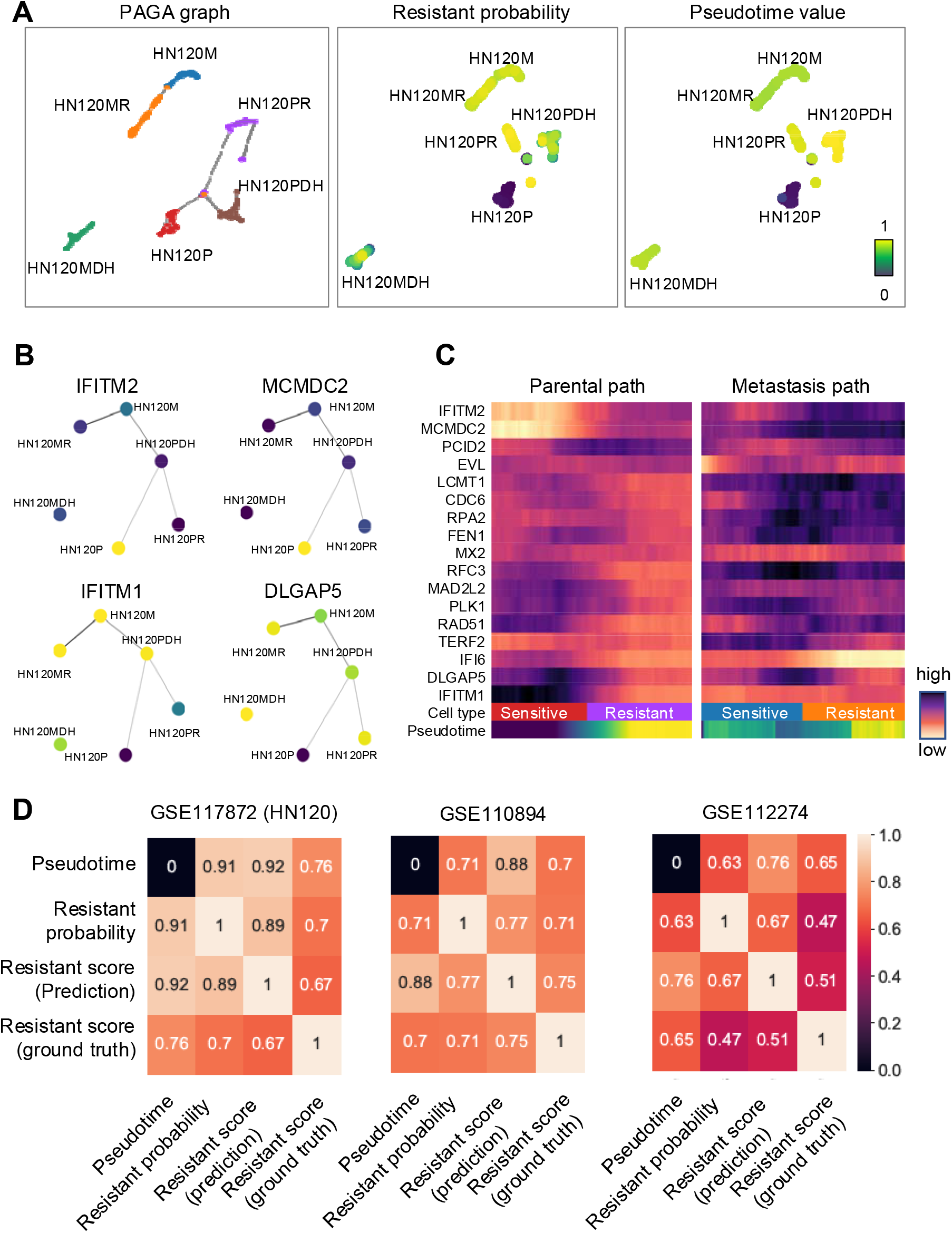
Connecting predicted drug response to pseudotime trajectory. **(A)** The diffusion map plot colored by annotated by six cell states annotation, predicted resistant probability, and diffusion pseudotime value on the Sharma (HN120) OSCC dataset. ‘P’ and ‘M’ stand for sensitive cells in ‘primary’ and ‘metastasis’ sites. ‘R’ and ‘DH’ stand for ‘resistant’ and ‘drug holiday’ cells. The combination of ‘P’/’M’ and “R’/’DH’ constructs six cell states. The Red **(B)** PAGA graph of the Sharma (HN120) dataset is colored by four critical genes. Nodes of the graph represent cell clusters annotated by six cell states. **(C)** The heatmap analysis towards the expression of critical genes in the sensitive-resistant state paths for parental cells and metastasis cells. Upper rows represent gene expressions, while columns represent cells. Cells are ordered by their response scores. **(D)** The correlation between diffusion pseudotime value and gene scores related to cell resistance across three datasets.

The PAGA graphs in **Fig. 4A** indicate strong connections between the diffusion pseudotime and predicted resistant probability in scDEAL. The diffusion pseudotime graph suggested setting the primary site of Sharma (HN120) cells as an origin root; its sensitivity towards Cisplatin decreases along with the cell trajectory. Beyond the consistency between prediction and the trajectory topology, we further explain the trend of resistance development by the critical genes identified by the integrate gradient method. Through the PAGA graphs (**Fig. 4B**), we observe that the expression levels of these genes are high within cells established from the primary site of the tumor and decrease elsewhere. On the contrary, expressions of resistant critical genes (such as IFITM and DLGAP5) are seen to increase towards the end of the sensitive–resistant path, respectively. We discovered consistent critical gene expression variation along with trajectory paths from sensitive to resistant, as shown in **Fig. 4C**. We observe changes of critical genes expressed in sensitive OSCC cells such as IFITM2 and MCMDC2. Similar trends are observed in three datasets, including Kong’s dataset (15) (GSE112274), Schnepp’s dataset (16) (GSE140440), and Sharma’s dataset (17) (GSE118782). We investigated three scores characterizing cell resistance in drug response, including the prediction probability for the resistant label, the resistance gene score derived from prediction, and the resistant score derived from the ground truth (**Fig. 4D**). Pearson’s correlations among three scores and the pseudotime value are commonly high, which indicated that predictions of scDEAL could highly imply drug resistance development.

## Discussion

Our work, for the first time, provides a novel method to augment single-cell data analyses and interpretation using bulk gene expression data, which can be widely applied to predict drug responses of cell populations in any cancer single-cell data. The neural networks adapted to single-cell data can be pre-trained on a large volume of bulk cell-line data. Hence, a large number of labeled single-cell data is not required to infer drug sensitivity prediction in our study. We benchmarked scDEAL on six scRNA-seq data with experimentally validated drug response labels, with an average F1 score of 0.78. For a prostate cancer dataset with 324 cells treated with the Docetaxel, scDEAL achieves the highest F1 as 0.97 and AUROC as 0.99. The robustness of scDEAL was demonstraed by performing random subsampling 1,000 times with a result of variance of F1 scores lower than 0.1. We also identified CGs corresponding to the Cisplatin responses in OSCC, showing distinct predicted response patterns in drug-sensitive and resistant cells. These CGs are regulated by TFs and enriched by pathways that have been reported to be related to the drug response. Our results also demonstrated the dynamic changes of drug sensitivities among cells and cell developmental trajectory.

The accuracy of the prediction results in scDEAL could vary, depending on the collection of bulk gene expression of cell lines. In the future, we will include more bulk-level data in training scDEAL. Moreover, databases, such as DrugCombDB (35), which includes 448,555 combinations of 2,887 drugs on 124 cell lines, can be included to train scDEAL for the prediction of combinatory drug responses in cell types. In addition, the genetic features between bulk and scRNA-seq data can be explained and biologically interpreted by the IG analysis. The predicted IGs can be used as targets for experimental validations on drug-gene relations via single-cell Perturb-seq (36). We believe our work can contribute and provide insights to cell reprogramming, drug selection and repurposing, and combinatory drug usage for improving therapeutic efficacy.

## Methods

### Datasets

We used the GDSC database as the source of bulk data which systematically collects 135,242 response profiles and genomic information across 565 compounds in 1,796 CCLs. GDSC is publicly available through the website (https://www.cancerrxgene.org/). Drug response annotation including half maximal inhibitory concentration (IC50) and area under the dose-response curve (AUC) are available through the page https://www.cancerrxgene.org/downloads/bulk_download. Gene expression data (RMA-normalized basal expression profiles) for cell lines can be access on GDSC (https://www.cancerrxgene.org/gdsc1000/GDSC1000_WebResources/Home.html).

All data analyzed within this paper are publicly available. The following data sets are available from the National Center for Biotechnology Information’s (NCBI) Gene Expression Omnibus (GEO, https://www.ncbi.nlm.nih.gov/geo/). All data are accessible through the GEO Series accession numbers: GSE117872 (Sharma, et al., Cisplatin treated oral squamous cell carcinoma), GSE112274 (Kong, et al., Gefitinib treated lung cancer), GSE110894 (Bell, et al., I-BET-762 treated acute myeloid leukemia), GSE140440 (Schnepp et al., Docetaxel treated prostate cancer), and GSE149383 (Aissa, et al., Erlotinib treated lung carcinoma) (**Table 1**).

**Table 1.** Data summary of the scRNA-seq datasets.

| Data | Drug | GEO access | Cells | Species | Cancer type | Ref |
| --- | --- | --- | --- | --- | --- | --- |
| Sharma, (HN120) | Cisplatin | GSE117872 | 548 | Homo sapiens | Oral squamous cell carcinomas | (17) |
| Sharma, (HN137) | Cisplatin | GSE117872 | 568 | Homo sapiens | Oral squamous cell carcinomas |  |
| Kong | Gefitinib | GSE112274 | 507 | Homo sapiens | Lung cancer | (15) |
| Aissa | Erlotinib | GSE149383 | 1496 | Mus musculus | Lung cancer | (37) |
| Bell | I-BET-762 | GSE110894 | 1419 | Mus musculus | Acute myeloid leukemia | (18) |
| Schnepp | Docetaxel | GSE140440 | 324 | Homo sapiens | Prostate Cancer | (16) |

### Pre-processing for expression profiles in GDSC dataset

In our experiment, we selected 1,018 cell lines and 17,419 genes from the expression profiles correspond to a subset of drug screening annotation for five drugs, including I-BET-762, Cisplatin, Gefitinib, Docetaxel, and Erlotinib, corresponding to the drug treatment in the single-cell data. All 1,018 cell expression profiles in GDSC are merged to their corresponding drug screening profiles according to their Catalogue of Somatic Mutations in Cancer (COSMIC) ID. Each cell line will be annotated with their drug screening records including AUC values, cell line names, and drug names. After the merging, there are 808 cell lines that have drug screening for our selected drugs. Expression values in the profiles are then scaled from 0 to 1 by “*pre-processing*.*MinMaxScaler*” in the package sklearn (38) before the training. The GDSC dataset is then split into training, validation, and test sets with the proportion of 60%, 20%, and 20%, respectively. The split is a stratified split where the proportion of sensitive and resistant cells are identical across training set, validation, and test set. The split is performed by the function “*train_test_split*” in the package sklearn.

### Pre-processing for labels transforming drug AUC to binary label in GDSC dataset

Our drug response labels of cell lines to each drug in the bulk-level gene are derived from the AUC values in the GDSC database. They are binarized with the waterfall method describe by the CCLE study (39) while cell lines have a response to a specific a selected drug is annotated as 1 and resistant as 0. The waterfall method firstly sorts cell lines according to their AUC values in descending order and generates an AUC-cell line curve in which the x-axis represents cell lines and the y-axis represents AUC values. The cutoff of AUC values is determined in two strategies: 1) for linear curves (whose regression line fitting has a Pearson correlation >0.95), the sensitive/resistant cutoff of AUC values is the median among all cell lines; 2) otherwise, the cut off is the AUC value of a specific boundary data point. It has the largest distance to a line linking two datapoints having the largest and smallest AUC values. Waterfall plots of drugs we applied in **scDEAL** are present in **(Supplementary Fig. S2)**.

### Data sampling for predictor training

As described in the label binarization, the proportion of sensitive and resistant cell lines are different across drug treatments **(Supplementary Fig. S3**). Potentially, the imbalance of drug response labels in the training set may affect the performance of the model. We, therefore, introduce different sampling methods as options to balance the proportion of sensitive and resistant cell lines when training the prediction model in the bulk level shown in. Three sampling methods as hyperparameters are introduced in the bulk model training, including up-sampling, down-sampling, and SMOTE-sampling (40). Up-sampling randomly duplicates samples in the minority, while down-sampling discards samples in the majority class to generate a training set that having the same number of sensitive and resistant cell lines, respectively. SMOTE balances the training set in a similar manner to up-sampling. It generates synthetic cell lines by selecting k nearest neighbors for a random sample in the minority class and synthesis a novel artificial sample within neighbors in feature space. All sampling methods are implemented by the python library imblearn (41).

### Pre-processing for scRNA-seq data

Quality control and pre-processing of the scRNA-seq data were performed using python package SCANPY (42). Specifically, cells with less than 200 detected genes (indicative of no cell in the droplet), and genes detected in less than 3 cells are filtered out using the function “*filter_cells*” and “*filter_genes*”. Percentages of mitochondrial genes expression and numbers of genes that have UMI count > 1 were calculated by the function “*cal_ncount_ngenes*”. Counts matrices were normalized dividing by the total UMI count in each cell, multiplied by a factor of 10,000 using the function “*normalize_total*”, and log one plus transformed using the function “*log1p*”. All scRNA-seq dataset are then split into training set, validation, and test set with the proportion of 60%, 20%, and 20% respectively. The split is a stratified split where the proportion of sensitive and resistant cells are identical across training, validation, and test sets. The split is performed by the function “*train_test_split*” in the package sklearn.

### scDEAL workflow design

#### Step 1. Bulk gene feature extractor

The first introduced step of transfer learning is the gene feature extractor. Gene feature extractors are applied to extract variable gene features, reduce data dimensionality and denoise data. Also, it is a fined-tuned pre-training step for the sensitivity prediction model P that has better initial model weights than random initialization. Our solutions of feature extractors are non-linear neural networks (NNs): Auto Encoder (43) (AE) and Variational Auto Encoders (44) (VAE).

The feature extractor *M*_*b*_ is modeled as an unsupervised autoencoder to learn a low dimensional representation from a bulk RNA-seq expression matrix *X*_*b*_, where each row of the matrix represents a cell and each column of the matrix is a gene. The basic architecture of an autoencoder is composed of two NNs: an encoder (*M*_*b*_) and a decoder (*DEC*_*b*_). *M*_*b*_ subtracts the data to a lower-dimensional subspace, and the *DEC*_*b*_ reconstructs the low dimensional representations back an approximation of *X*_*b*_ . Parameters inside the autoencoder are optimized by the reconstruction loss function (Mean Square Error, MSE) between the output of *DEC*_*b*_ and *X*_*b*_, aiming to make the reconstructed matrix as similar as *X*_*b*_. The model can be trained by equation *Eq. 1* as follows:

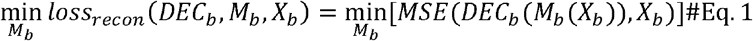

Alternatively, the feature extractor *M*_*b*_ can also be modeled as an unsupervised Variational Autoencoder (VAE) (44) to learn a low dimensional representation from *X*_*b*_ . Like AE, the architecture of a VAE is also composed of an encoder (*X*_*b*_) and a decoder (*DEC*_*b*_). However, VAE is a generative model, which applies NNs to estimate parameters of a latent distribution that enable us to sample data in its lower-dimensional representation. In this model, an encoder (*M*_*b*_) is applied to estimate the mean (μ) and variance (σ) of the data distribution in the lower-dimensional space. The decoder (*DEC*_*b*_) received the low dimensional sampling result to generate an approximation of *X*_*b*_ . Parameters inside the VAE are optimized by the weighted sum of the reconstruction loss function between the output of *DEC*_*b*_ and *X*_*b*_, and a Kullback–Leibler Divergence (KL), named the lower bound of likelihood loss (*loss*_*ELBO*_). The model can be trained by minimizing the loss explained by equation *Eq. 2* as follows:

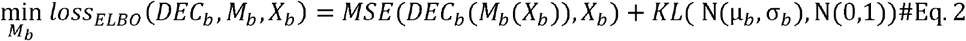

where N(µ,σ) is a normal distribution with mean µ and variance σ; *KL*() is the Kullback– Leibler Divergence between distributions. All trained parameters in Step 1 are listed in **Supplementary Table S3**.

#### Step 2. Single-cell gene feature extractor

The second step of the transfer learning is the d single-cell gene feature extractor. It is also applied to feature extractors are applied to extract low-dimensional gene features. Besides, it is a fined-tuned pre-training step for the transfer learning model in the single-cell level in Step 5. Its model structure, training process, and loss function are identical to the previously introduced feature extractor at the bulk level. However, its input and reconstruction target are substituted by the scRNA-seq data *X*_*s*_. All trained parameters in Step 2 are listed in **Supplementary Table S4**.

#### Step 3. Bulk drug response predictor

The third introduced step of transfer learning is the drug response predictor. We pre-train a NN model (*P*) based on a fully connected multi-layer perceptron (MLP) (45) on bulk RNA-seq data to estimate the correlation of drug response and bulk gene expressions. Parameters inside *P* are optimized with the classification loss (i.e., cross-entropy) between the predictive value of sensitivity *DEC*_*b*_ (*M*_*b*_(*X*_*b*_)) and the ground truth *Y*_*b*_, which can be trained by:

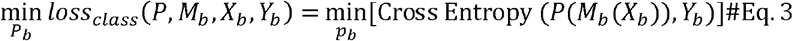

The result of Step 2 (P), will be finally applied to link the level of drug response cells of*M*_*s*_ cell-drug in Step 5.

#### Step 4. Transfer learning

In this step, we will introduce the transfer learning model, which is the core step of the model. It adapts the gene features extracted from bulk and single level to enable the sensitivity prediction for cells through the predictor P.

We apply a DaNN (46) model to derive the feature extractor *M*_*s*_ in the single-cell level. The DaNN model introduces an extra loss named the Maximum Mean Discrepancy (MMD) to estimate the similarity of output *M*_*b*_ and *M*_*s*_. This similarity between two gene features is added to the classification loss during the training process of the predictor *P* to ensure that the feature space from the output of *M*_*s*_ exhibits similar distributions to the output of *M*_*B*_. This transfer learning model (D) takes two sources of data *X*_*b*_ and *X*_*s*_ as inputs, and *Y*_*s*_ as output. DaNN models are trained to update two gene extractors *M*_*B*_ and *M*_*s*_ at the same time. The loss of DaNN marked as *loss*_*DaNN*_ can be interpreted as a weighted sum of the bulk level prediction loss *loss*_*class*_, and the gene feature distribution *loss*_*MMD*_. *loss*_*class*_ is the classification loss of the sensitivity prediction of bulk RNA-seq data, which is the loss described in equation *Eq. 3*. The distribution loss *loss*_*MMD*_ is the discrepancy between the feature output of bulk and single-cell data. The *loss*_*MMD*_ function is described in *Eq. 4*:

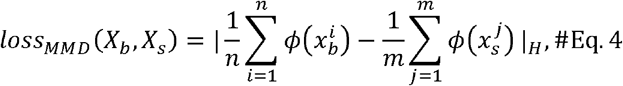

where 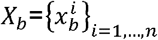 and 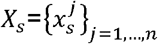 are data vectors for bulk and single-cell data; ϕ (.) is referred to as the feature space map to the universal Reproducing Kernel Hilbert Space (RKHS). The RKHS norm | · |_*H*_ is applied to measure the distance of two vectors with different dimensions. At last, the overall loss of DaNN is defined as

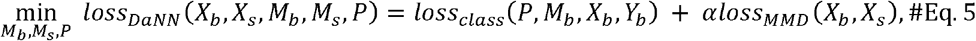

where α is a weight of *loss*_*MMD*_ .Minimizing the transfer learning loss *loss*_*DaNN*_, the result will be the extracted and matched gene features output from *M*_*s*_ and will be used in the sensitivity prediction in Step 5.

#### Single-cell Prediction

The last step of transfer learning framework is to generate drug response prediction for cells. The trained *M*_*s*_ in Step 4 is a model after feature transformation. Therefore, the gene feature from scRNA-seq data extracted by *M*_*s*_ can be recognized by the predictor *P*. Therefore, we will ensemble two components *M*_*s*_ and *P* as a drug response prediction network at the single-cell level. The prediction network will take scRNA-seq data *X*_*s*_ as input and output binary drug response (sensitive or resistant) for each cell in *X*_*s*_ .The binary drug response predictions will be applied to calculate a sensitive/resistant score based on the expression level of marker genes in the predicted sensitive/resistant cluster. The detailed method is described in the **Sensitive and resistant gene score analysis** section.

### Benchmarking metrics for the scDEAL prediction

To evaluate the prediction of scDEAL, we applied three metrics F-score, AUROC, and AP which are measurements of a classification test’s accuracy.

***F-score*** can be interpreted as a weighted average of precision and recall. F-score reaches its highest value at 1 and lowest score at 0. The equation for the F-score is:

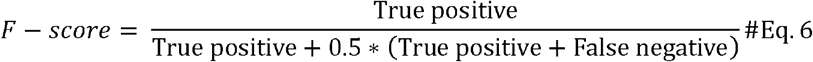

We implemented F-score tests using the *“f1_score*” function in the package sklearn (38).

***AUROC score*** computes the area under the receiver operating characteristic (ROC) curve. ROC curve’s x-axis is the true positive rate and the y-axis is the false positive rate derived from prediction scores. The curve is generated by setting different thresholds to binarize the numerical prediction scores. AUROC computes the area under the precision-recall curve with the trapezoidal rule, which uses linear interpolation. We implemented AUROC tests using the *“roc_auc_score*” function in the sklearn package.

***AP score*** summarizes a precision-recall curve (PRC) as the weighted mean of precisions achieved at each threshold, with the increase in recall from the previous threshold used as the weight. The equation of AP score is:

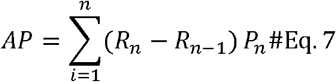

where *P*_*n*_ and *R*_*n*_ are the precision and recall at a threshold *n* ordered by its value. We implemented AP tests using the “*average_precision_score*” function in the package sklearn.

### Sensitive and resistant gene score analysis

To define the sensitive and resistant score, we collected differentially expressed genes between the sensitive and the resistant cluster based on both the predictions and annotations in the dataset (**Table 1**). The sensitive or resistant gene score is derived from the binned average expression of a set of differentially expressed genes either in the sensitive or resistant cell cluster. The score of each set of genes was evaluated with the *“score_genes”* function built in the SCANPY (42). To select the differentially expressed genes, we performed Wilcoxon signed-rank tests using the “*rank_genes_groups*” function. By default, the top 50 genes (whose adjusted p-values<0.05) ranked by the z-score are selected to calculate the sensitive or resistant gene score.

### Data sampling and repetition for stability tests

The stability test is performed by retraining the DTL model ten times with different random seeds and randomly sampled subsets of cells. To preserve the original sensitive and resistant cell ratio in the dataset, we choose to perform the stratified sampling of the sensitive and resistant cells in the dataset. The sampling is performed using the “*resample*” function in the **sklearn** package (38). The number of output (n_samples parameter) is set to be 80% of the input, and the sampled data points cannot be sampled again with the setting “replace=False”.

### Critical gene identification with Integrated Gradients

We applied integrated gradients (**IG**) (47) to characterize critical input genes features in the scDEAL model. IG represents the integral of gradients with respect to each gene expression as inputs along the path from zero expression as a baseline to the input expression level (**Supplementary Fig. S4**). The integral is approximated using the Riemann rule described as follows:

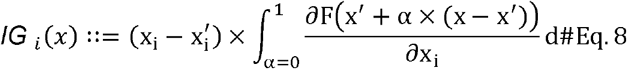

It calculated the importance of the i-th gene expression of the input cell x. αis the scaling coefficient; 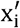 is the baseline expression level gene i which is 0 in our case; and ∂F(x) / ∂x_i_ represents the gradient of F(x) along the i -th dimension.

We apply the “*IntegratedGradients*” class in the python Captum library (48) to calculate IG values. The input is the function is our expression matrix, trained model, and output labels. The output of the function is IG matrices of the same shape as the input expression matrix. Rows resents genes and columns represent cells. Values in a matrix are the corresponding IG values.

Since scDEAL is a binary classification deep learning model. It has two nodes in the output layer to predict sensitive and resistant probability. Based on the sensitivity or resistance for each gene contribute, we can obtain two IG matrices for each input data with corresponds to the sensitive and resistant output, respectively. The IG matrix can be found in the model output file “*attr_integrated_gradient*.*h5ad*”. The IG matrix is stored with an “*AnnData*” object and can be read by the function “*sc*.*read_h5ad*”.

To select critical genes that have significantly higher IG values within the sensitive (or resistant) cell cluster, we utilize the Wilcoxon test using the function “*sc*.*tl*.*rank_genes_groups*” between Cisplatin sensitive and resistant cells in SCANPY. We consider the genes which have Bonferroni adjusted p-values less than 0.05 and log-fold changes greater than 1 as acritical genes. Sensitive critical genes and resistant critical genes are further determined depending on the predicted sensitive and resistant cluster labels.

### Functional enrichment

The KEGG and GO gene set enrichment analysis is performed using R package **ClusterProfiler** (49). The gene symbols are first converted to ENTREZID by the “*bitr*” function. The KEGG analysis is performed via the function “*enrichKEGG*” with the parameter setting “*qvalueCutoff*=0.05”. The GO analysis is performed by utilizing the function “*enrichGO*” with the parameter setting “*ont* = “ALL”, *pAdjustMethod* = “BH”, *qvalueCutoff* = 0.05”. GO biological process enrichment analysis was performed using the ClusterProfiler R package with a p-value cut-off of 0.01 and a q-value cut-off of 0.05.

### TF and Motif analysis

Epigenetic Landscape in Silico deletion Analysis (**LISA**) (27) was used to identify potential transcription factors (TFs) and chromatin regulators (CRs) that regulate the sensitive and resistant critical genes, respectively. LISA first models the epigenetic landscape based on the input marker genes as well as public epigenomic profiles (DNase-seq, H3K27ac ChIP-seq) in CistromeDB, then performs in silico detection of TF binding sites on the epigenetic landscape to evaluate the essentiality of the transcriptional regulators. We utilize the default setting of LISA2 with the parameter *‘hg38’, rp_map = ‘enhanced_10K’, assays = [‘Direct’,’DNase’,’H3K27ac’], isd_method = ‘chipseq’, verbose = 1)*. The number of background genes is set for 3000. The motifs of TFs in LISA2 are fetched by JASPAR (50) database.

### Trajectory inference for Oral Squamous Cell Carcinomas

The trajectory analysis is carried out in the python package SCANPY (42). The PCA and nearest neighbor graph required by the trajectory inference are generated in the step of analysis for scRNA-seq gene expression. We apply ForceAtlas2 (51) layout to preserves a graph topology that is better to be visualized using function “*draw_grap*.” To denoise the graph, we applied diffusion map (34) using the function “*diffmap*” to computing distances based on diffusion components similar to denoising a data matrix using PCA. The trajectory is then mapped to a backbone PAGA (33) connectivity structure using the “*paga*” function. By quantifying the connectivity cell types in the single-cell graph, PAGA generates a simplified graph of partitions, in which edge weights represent confidence in the presence of connections.

## Code availability

The source code of scDEAL is freely available on (https://github.com/OSU-BMBL/scDEAL).

## Authors’ Contributions

Q.M. conceived the basic idea and designed the framework. J.C., R.Q., Z.W., A.M., and J.Z. led framework construction and performed data analysis. D.X. helped the DLT model optimization, and L.L. provided valuable comments for the interpretation of drug responses. All authors participated in the interpretation and writing the manuscript.

## Acknowledgements

This work was supported by funding from the National Institutes of Health (R01-131399 and R35-GM126985). The project described was also supported by Award Number Grant UL1TR002733 from the National Center For Advancing Translational Sciences. The authors would like to thank Dr. Fei He (Northeast Normal University, China) for his assistance and advice in the pipeline construction.

## References

1. Verjans, E.T., Doijen, J., Luyten, W., Landuyt, B. and Schoofs, L. (2018) Three□dimensional cell culture models for anticancer drug screening: Worth the effort? Journal of cellular physiology, 233, 2993–3003.

2. Schirle, M. and Jenkins, J.L. (2016) Identifying compound efficacy targets in phenotypic drug discovery. Drug discovery today, 21, 82–89.

3. Wong, C.H., Siah, K.W. and Lo, A.W. (2019) Estimation of clinical trial success rates and related parameters. Biostatistics, 20, 273–286.

4. Rambow, F., Rogiers, A., Marin-Bejar, O., Aibar, S., Femel, J., Dewaele, M., Karras, P., Brown, D., Chang, Y.H., Debiec-Rychter, M. et al.. (2018) Toward Minimal Residual Disease-Directed Therapy in Melanoma. Cell, 174, 843–855 e819.

5. Wang, J., Ma, A., Chang, Y., Gong, J., Jiang, Y., Qi, R., Wang, C., Fu, H., Ma, Q. and Xu, D. (2021) scGNN is a novel graph neural network framework for single-cell RNA-Seq analyses. Nature Communications, 12, 1882.

6. Gayoso, A., Steier, Z., Lopez, R., Regier, J., Nazor, K.L., Streets, A. and Yosef, N. (2021) Joint probabilistic modeling of single-cell multi-omic data with totalVI. Nature Methods, 18, 272–282.

7. Wang, J., Agarwal, D., Huang, M., Hu, G., Zhou, Z., Ye, C. and Zhang, N.R. (2019) Data denoising with transfer learning in single-cell transcriptomics. Nature methods, 16, 875–878.

8. Wu, Z., Lawrence, P.J., Ma, A., Zhu, J., Xu, D. and Ma, Q. (2020) Single-Cell Techniques and Deep Learning in Predicting Drug Response. Trends in Pharmacological Sciences, 41, 1050–1065.

9. Tan, C., Sun, F., Kong, T., Zhang, W., Yang, C. and Liu, C. (2018), International Conference on Artificial Neural Networks. Springer, pp. 270–279.

10. Dhruba, S.R., Rahman, R., Matlock, K., Ghosh, S. and Pal, R. (2018) Application of transfer learning for cancer drug sensitivity prediction. BMC Bioinformatics, 19, 497.

11. Pan, S.J. and Yang, Q. (2009) A survey on transfer learning. IEEE Transactions on knowledge and data engineering, 22, 1345–1359.

12. Yosinski, J., Clune, J., Bengio, Y. and Lipson, H. (2014) How transferable are features in deep neural networks? arXiv preprint 1411.1792.

13. Yang, W., Soares, J., Greninger, P., Edelman, E.J., Lightfoot, H., Forbes, S., Bindal, N., Beare, D., Smith, J.A. and Thompson, I.R. (2012) Genomics of Drug Sensitivity in Cancer (GDSC): a resource for therapeutic biomarker discovery in cancer cells. Nucleic acids research, 41, D955–D961.

14. Iorio, F., Knijnenburg, T.A., Vis, D.J., Bignell, G.R., Menden, M.P., Schubert, M., Aben, N., Gonçalves, E., Barthorpe, S. and Lightfoot, H. (2016) A landscape of pharmacogenomic interactions in cancer. Cell, 166, 740–754.

15. Kong, S.L., Li, H., Tai, J.A., Courtois, E.T., Poh, H.M., Lau, D.P., Haw, Y.X., Iyer, N.G., Tan, D.S.W. and Prabhakar, S. (2019) Concurrent Single-Cell RNA and Targeted DNA Sequencing on an Automated Platform for Comeasurement of Genomic and Transcriptomic Signatures. Clinical chemistry, 65, 272–281.

16. Schnepp, P.M., Shelley, G., Dai, J., Wakim, N., Jiang, H., Mizokami, A. and Keller, E.T. (2020) Single-Cell Transcriptomics Analysis Identifies Nuclear Protein 1 as a Regulator of Docetaxel Resistance in Prostate Cancer Cells. Molecular Cancer Research, 18, 1290–1301.

17. Sharma, A., Cao, E.Y., Kumar, V., Zhang, X., Leong, H.S., Wong, A.M.L., Ramakrishnan, N., Hakimullah, M., Teo, H.M.V. and Chong, F.T. (2018) Longitudinal single-cell RNA sequencing of patient-derived primary cells reveals drug-induced infidelity in stem cell hierarchy. Nature communications, 9, 4931.

18. Bell, C.C., Fennell, K.A., Chan, Y.-C., Rambow, F., Yeung, M.M., Vassiliadis, D., Lara, L., Yeh, P., Martelotto, L.G. and Rogiers, A. (2019) Targeting enhancer switching overcomes non-genetic drug resistance in acute myeloid leukaemia. Nature communications, 10, 1–15.

19. Ghosh, S. (2019) Cisplatin: The first metal based anticancer drug. Bioorganic chemistry, 88, 102925.

20. Dasari, S. and Tchounwou, P.B. (2014) Cisplatin in cancer therapy: molecular mechanisms of action. European journal of pharmacology, 740, 364–378.

21. Zhai, Y., Li, N., Jiang, H., Huang, X., Gao, N. and Tye, B.K. (2017) Unique Roles of the Non-identical MCM Subunits in DNA Replication Licensing. Mol Cell, 67, 168–179.

22. Naiki, T., Shimomura, T., Kondo, T., Matsumoto, K. and Sugimoto, K. (2000) Rfc5, in cooperation with rad24, controls DNA damage checkpoints throughout the cell cycle in Saccharomyces cerevisiae. Mol Cell Biol, 20, 5888–5896.

23. Kim, S.H., Ho, J.N., Jin, H., Lee, S.C., Lee, S.E., Hong, S.K., Lee, J.W., Lee, E.S. and Byun, S.S. (2016) Upregulated expression of BCL2, MCM7, and CCNE1 indicate cisplatin-resistance in the set of two human bladder cancer cell lines: T24 cisplatin sensitive and T24R2 cisplatin resistant bladder cancer cell lines. Investig Clin Urol, 57, 63–72.

24. Yánez, D.C., Ross, S. and Crompton, T. (2020) The IFITM protein family in adaptive immunity. Immunology, 159, 365–372.

25. John, D.S., Aschenbach, J., Krüger, B., Sendler, M., Weiss, F.U., Mayerle, J., Lerch, M.M. and Aghdassi, A.A. (2019) Deficiency of cathepsin C ameliorates severity of acute pancreatitis by reduction of neutrophil elastase activation and cleavage of Ecadherin. J Biol Chem, 294, 697–707.

26. Khaket, T.P., Singh, M.P., Khan, I. and Kang, S.C. (2020) In vitro and in vivo studies on potentiation of curcumin-induced lysosomal-dependent apoptosis upon silencing of cathepsin C in colorectal cancer cells. Pharmacol Res, 161, 105156.

27. Qin, Q., Fan, J., Zheng, R., Wan, C., Mei, S., Wu, Q., Sun, H., Brown, M., Zhang, J. and Meyer, C.A. (2020) Lisa: inferring transcriptional regulators through integrative modeling of public chromatin accessibility and ChIP-seq data. Genome biology, 21, 1–14.

28. Funato, T., Kozawa, K., Kaku, M. and Sasaki, T. (2001) Modification of the sensitivity to cisplatin with c-myc over-expression or down-regulation in colon cancer cells. Anti-cancer drugs, 12, 829–834.

29. Makovec, T. (2019) Cisplatin and beyond: molecular mechanisms of action and drug resistance development in cancer chemotherapy. Radiology and oncology, 53, 148–158.

30. Galluzzi, L., Senovilla, L., Vitale, I., Michels, J., Martins, I., Kepp, O., Castedo, M. and Kroemer, G. (2012) Molecular mechanisms of cisplatin resistance. Oncogene, 31, 1869–1883.

31. Galluzzi, L., Vitale, I., Michels, J., Brenner, C., Szabadkai, G., Harel-Bellan, A., Castedo, M. and Kroemer, G. (2014) Systems biology of cisplatin resistance: past, present and future. Cell death & disease, 5, e1257–e1257.

32. Kimura, A., Ishida, Y., Inagaki, M., Nakamura, Y., Sanke, T., Mukaida, N. and Kondo, T. (2012) Interferon-γ is protective in cisplatin-induced renal injury by enhancing autophagic flux. Kidney international, 82, 1093–1104.

33. Wolf, F.A., Hamey, F.K., Plass, M., Solana, J., Dahlin, J.S., Göttgens, B., Rajewsky, N., Simon, L. and Theis, F.J. (2019) PAGA: graph abstraction reconciles clustering with trajectory inference through a topology preserving map of single cells. Genome biology, 20, 59.

34. Haghverdi, L., Büttner, M., Wolf, F.A., Buettner, F. and Theis, F.J. (2016) Diffusion pseudotime robustly reconstructs lineage branching. Nature methods, 13, 845.

35. Liu, H., Zhang, W., Zou, B., Wang, J., Deng, Y. and Deng, L. (2019) DrugCombDB: a comprehensive database of drug combinations toward the discovery of combinatorial therapy. Nucleic Acids Research, 48, D871–D881.

36. Dixit, A., Parnas, O., Li, B., Chen, J., Fulco, C.P., Jerby-Arnon, L., Marjanovic, N.D., Dionne, D., Burks, T., Raychowdhury, R. et al.. (2016) Perturb-Seq: Dissecting Molecular Circuits with Scalable Single-Cell RNA Profiling of Pooled Genetic Screens. Cell, 167, 1853–1866 e1817.

37. Aissa, A.F., Islam, A., Ariss, M.M., Go, C.C., Rader, A.E., Conrardy, R.D., Gajda, A.M., Rubio-Perez, C., Valyi-Nagy, K., Pasquinelli, M. et al.. (2021) Single-cell transcriptional changes associated with drug tolerance and response to combination therapies in cancer. Nat Commun, 12, 1628.

38. Pedregosa, F., Varoquaux, G., Gramfort, A., Michel, V., Thirion, B., Grisel, O., Blondel, M., Prettenhofer, P., Weiss, R. and Dubourg, V. (2011) Scikit-learn: Machine learning in Python. the Journal of machine Learning research 12.

39. Barretina, J., Caponigro, G., Stransky, N., Venkatesan, K., Margolin, A.A., Kim, S., Wilson, C.J., Lehár, J., Kryukov, G.V. and Sonkin, D. (2012) The Cancer Cell Line Encyclopedia enables predictive modelling of anticancer drug sensitivity. Nature, 483, 603.

40. Chawla, N.V., Bowyer, K.W., Hall, L.O. and Kegelmeyer, W.P. (2002) SMOTE: synthetic minority over-sampling technique. Journal of artificial intelligence research, 16, 321–357.

41. Lemaître, G., Nogueira, F. and Aridas, C.K. (2017) Imbalanced-learn: A python toolbox to tackle the curse of imbalanced datasets in machine learning. The Journal of Machine Learning Research, 18, 559–563.

42. Wolf, F.A., Angerer, P. and Theis, F.J. (2018) SCANPY: large-scale single-cell gene expression data analysis. Genome biology, 19, 1–5.

43. Kramer, M.A. (1991) Nonlinear principal component analysis using autoassociative neural networks. AIChE journal, 37, 233–243.

44. Kingma, D.P. and Welling, M. (2013) Auto-encoding variational bayes. arXiv preprint 1312.6114.

45. Gardner, M.W. and Dorling, S. (1998) Artificial neural networks (the multilayer perceptron)—a review of applications in the atmospheric sciences. Atmospheric environment, 32, 2627–2636.

46. Ghifary, M., Kleijn, W.B. and Zhang, M. (2014), Pacific Rim international conference on artificial intelligence. Springer, pp. 898–904.

47. Sundararajan, M., Taly, A. and Yan, Q. (2017), International Conference on Machine Learning. PMLR, pp. 3319–3328.

48. Kokhlikyan, N., Miglani, V., Martin, M., Wang, E., Alsallakh, B., Reynolds, J., Melnikov, A., Kliushkina, N., Araya, C. and Yan, S. (2020) Captum: A unified and generic model interpretability library for pytorch. arXiv preprint 2009.07896.

49. Yu, G., Wang, L.-G., Han, Y. and He, Q.-Y. (2012) clusterProfiler: an R Package for Comparing Biological Themes Among Gene Clusters. OMICS: A Journal of Integrative Biology, 16, 284–287.

50. Fornes, O., Castro-Mondragon, J.A., Khan, A., van der Lee, R., Zhang, X., Richmond, P.A., Modi, B.P., Correard, S., Gheorghe, M., Baranašić, D. et al.. (2019) JASPAR 2020: update of the open-access database of transcription factor binding profiles. Nucleic Acids Research, 48, D87–D92.

51. Jacomy, M., Venturini, T., Heymann, S. and Bastian, M. (2014) ForceAtlas2, a continuous graph layout algorithm for handy network visualization designed for the Gephi software. PloS one, 9, e98679.

