## Supplementary Figures and Tables for "Deep Transfer Learning of Drug Responses by Integrating Bulk and Single-cell RNA-seq data"

Supplementary Table S1. Classification report for source models.

Supplementary Table S2. Stability of prediction results.

Supplementary Table S3. Parameters in training autoencoder for bulk feature extraction.

Supplementary Table S4. Parameters in training autoencoder for single-cell feature extraction.

Supplementary Fig. S1 Critical gene and pathways.

Supplementary Fig. S3. RPC/ROC plots for source models of different drugs.

Supplementary Fig. S2. Waterfall plots and min-max distance plots of the binarization on drug response profiles of all cell lines for nine selected drugs from the bulk RNA-Seq data of GDSC dataset.

Supplementary Fig. S4. Procedure of the Integrated Gradient method on the gene interpretation from the DTL model.

Supplementary Table S1. Classification report for source models.

| **GDSC drugs** | **Precision** | **Recall** | **F1-score** | **AUROC score** | **AP score** |
| --- | --- | --- | --- | --- | --- |
| I-BET-762 | 0.8424 | 0.8447 | 0.8435 | 0.7656 | 0.4891 |
| Cisplatin_64 | 0.8623 | 0.8882 | 0.8582 | 0.7446 | 0.4289 |
| Cisplatin_128 | 0.8891 | 0.9006 | 0.8923 | 0.7876 | 0.4802 |
| Gefitinib | 0.8367 | 0.8509 | 0.8431 | 0.7604 | 0.45 |
| Docetaxel | 0.6562 | 0.6584 | 0.6552 | 0.7097 | 0.6546 |
| Erlotinib | 0.8316 | 0.8198 | 0.8255 | 0.7046 | 0.3228 |

Supplementary Table S2. Stability of prediction results.

| **Group** | **Metric** | **Mean** | **Standard deviation** |
| --- | --- | --- | --- |
| Repetition | AP | 0.8691 | 0.0225 |
| Repetition | AUROC | 0.9143 | 0.0168 |
| Repetition | F1 | 0.8340 | 0.0224 |
| Repetition | Pearson correlation  (Resistant score) | 0.9750 | 0.0099 |
| Repetition | Pearson correlation  (Sensitive score) | 0.9254 | 0.0272 |
| Stratified sampling | AP | 0.8745 | 0.0338 |
| Stratified sampling | AUROC | 0.9171 | 0.0142 |
| Stratified sampling | F1 | 0.8493 | 0.0293 |
| Stratified sampling | Pearson correlation  (Resistant score) | 0.7456 | 0.0227 |
| Stratified sampling | Pearson correlation  (Sensitive score) | 0.8652 | 0.0079 |

Supplementary Table S3. Parameters in training autoencoder for bulk feature extraction. Abbreviations: Autoencoder (AE); Variational autoencoder (VAE).

| **Drugs** | **Sampling** | **Bottle-neck** | **Encoder** | **Encoder**  **hidden dimensions** | **Predictor hidden**  **dimension** |
| --- | --- | --- | --- | --- | --- |
| I-BET-762 | Down-sampling | 256 | AE | (256, 256) | (128, 64) |
| Cisplatin | Up-sampling | 64 | VAE | (1024, 256) | (32, 16) |
| Cisplatin | Up-sampling | 128 | VAE | (512, 256) | (64, 32) |
| Gefitinib | Down-sampling | 256 | AE | (256, 256) | (128, 64) |
| Docetaxel | SMOTE-sampling | 100 | AE | (1024, 512, 256) | (64, 32) |
| Erlotinib | Downs-sampling | 128 | AE | (512, 256) | (64, 32) |

Supplementary Table S4. Parameters in training autoencoder for single-cell feature extraction.

| **Dataset** | **Bottle-neck** | **Encoder** | **Input features** | **Encoder**  **Hidden dimensions** | **Predictor**  **Hidden dimensions** |
| --- | --- | --- | --- | --- | --- |
| GSE110894 | 256 | AE | 6644 | (256, 256) | (128, 64) |
| GSE117872  HN120 | 64 | VAE | 7107 | (1024, 256) | (32, 16) |
| GSE117872  HN137 | 128 | VAE | 7107 | (512, 256) | (64, 32) |
| GSE112274 | 256 | AE | 13,065 | (256, 256) | (128, 64) |
| GSE140440 | 100 | AE | 14,559 | (1024, 512, 256) | (64, 32) |
| GSE149383 | 128 | AE | 6898 | (512, 256) | (64, 32) |

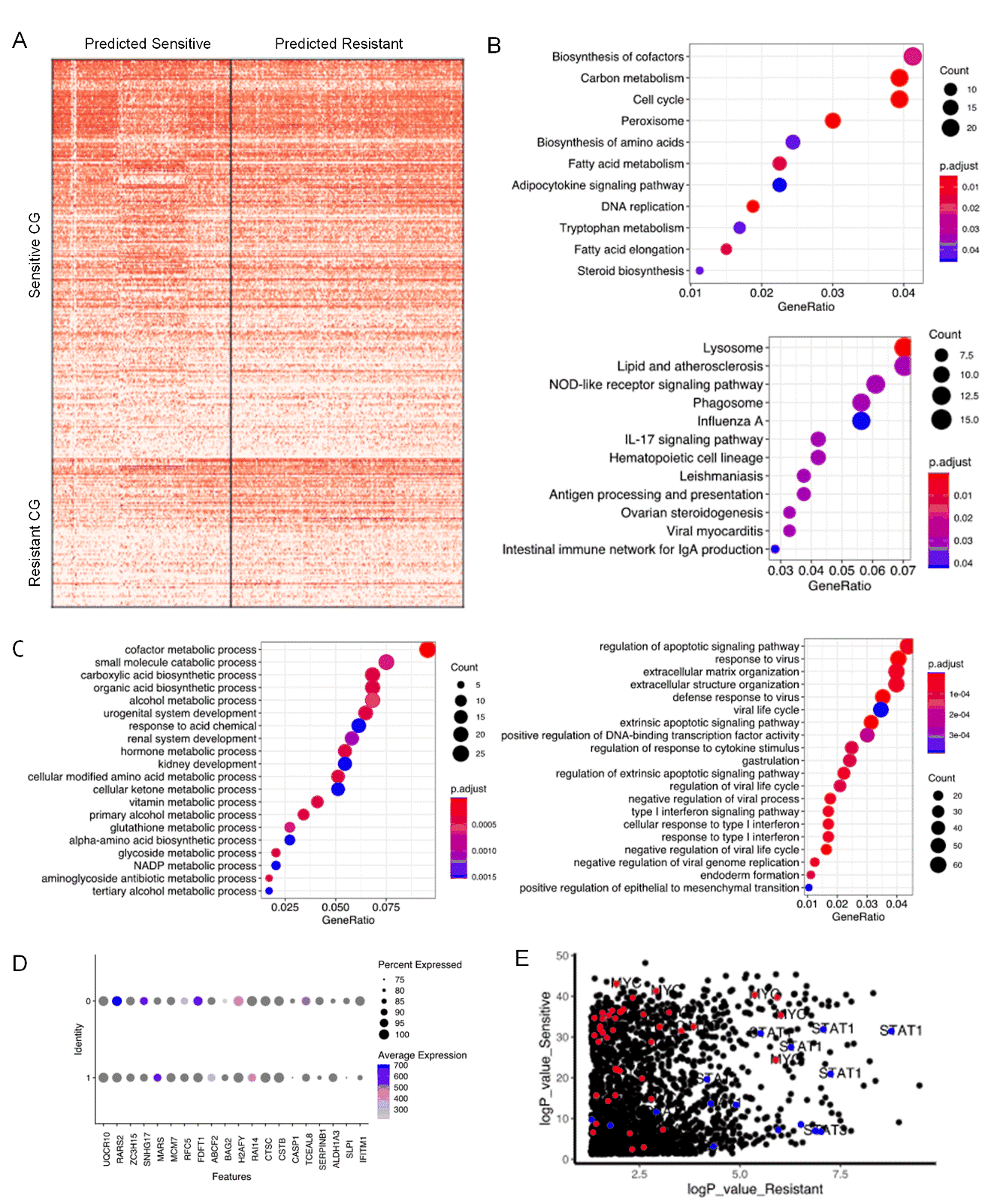

Supplementary Fig. S1 Critical gene and pathways. (A) heatmap of the expression of sensitive and resistant critical genes in two clusters. (B) KEGG analysis of sensitive critical genes (upper panel) and resistant critical genes (lower panel). (C) The GO analysis of differential expressed gene. The left panel shows the GO results for genes upregulated in the sensitive cluster, while the right panel shows the GO results for genes upregulated in the resistant cluster. (D) The expression of Top10 resistant and sensitive critical genes. (E) The identified regulatory TFs of sensitive CGs (red) and resistant CGs (blue).

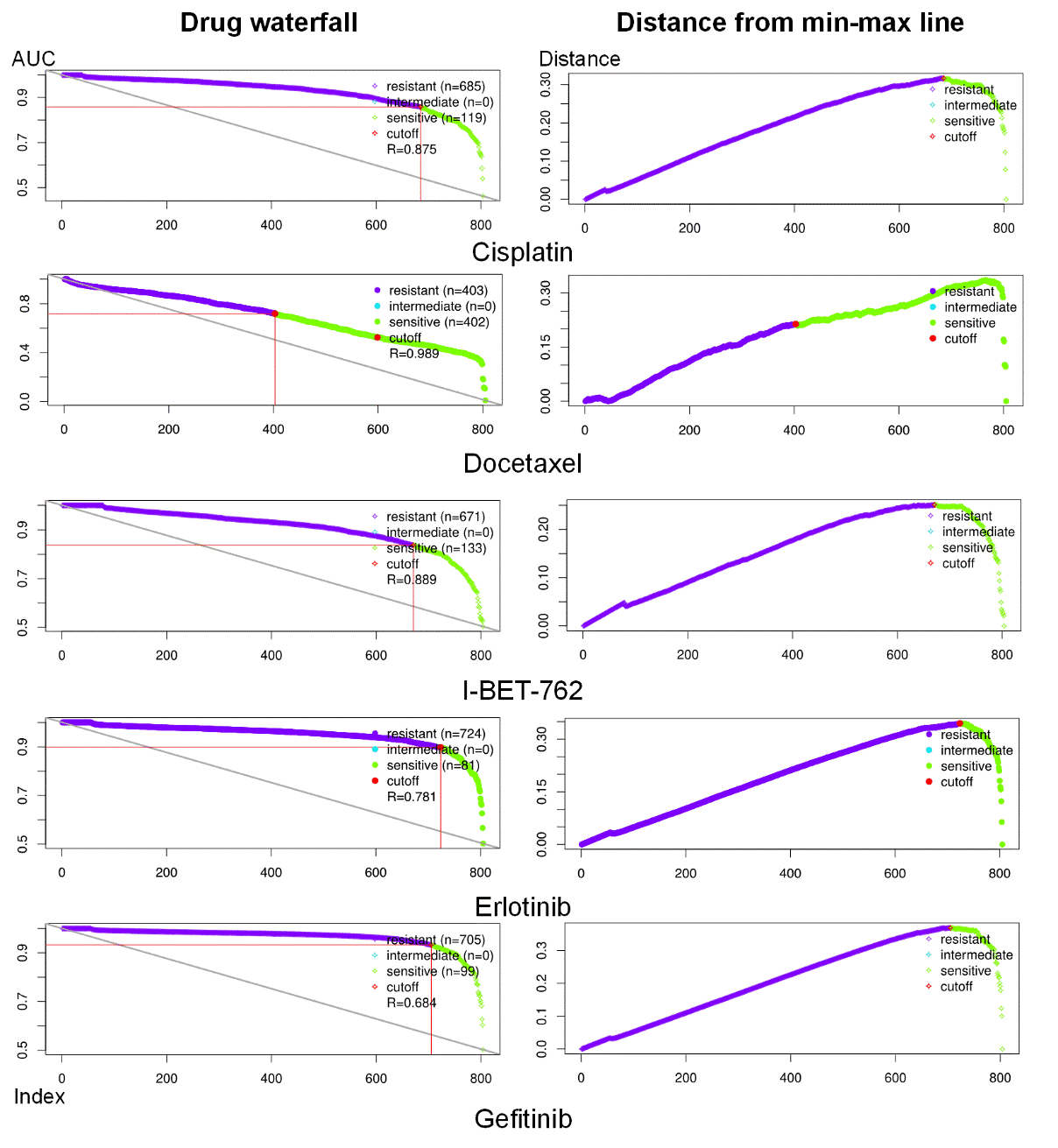

Supplementary Fig. S2. Waterfall plots and min-max distance plots of the binarization on drug response profiles of all cell lines for nine selected drugs from the bulk RNA-Seq data of GDSC dataset. The X-axis of each waterfall plot is the rank of cell line AUC values in descending order. The Y-axis of the upper panel is the AUC value, and the Y-axis in the lower panel is the distance to the line linking cell lines having the largest and the lowest AUC values. The red dots represent the cutoff of AUC values. Cell lines having AUC lower than the cutoff are marked as resistant and vice versa.

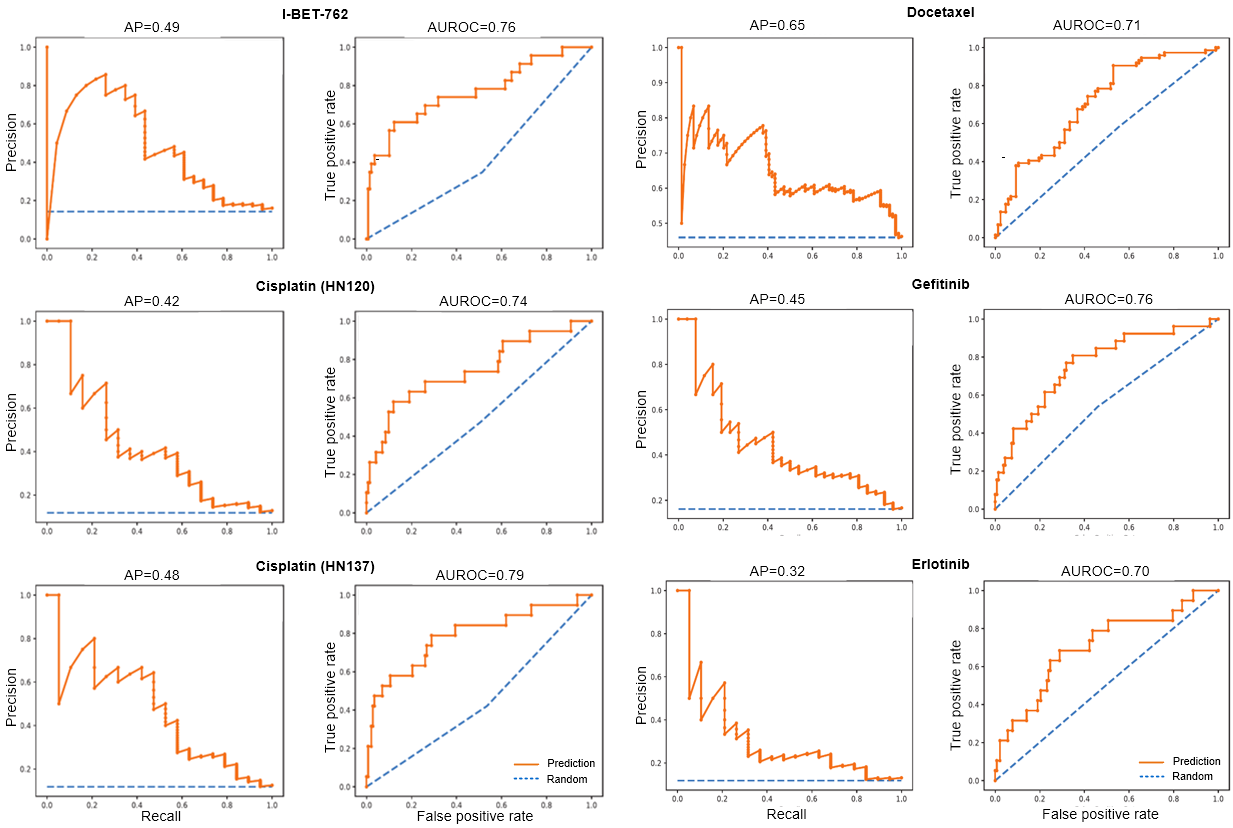

Supplementary Fig. S3. RPC/ROC plots for source models of different drugs. Curves on the left size are PRC curves while curves on the right side are PRC curves. Orange curves represent our prediction while blue lines are random predictions. Scores upon figures are their AP/AUROC scores.

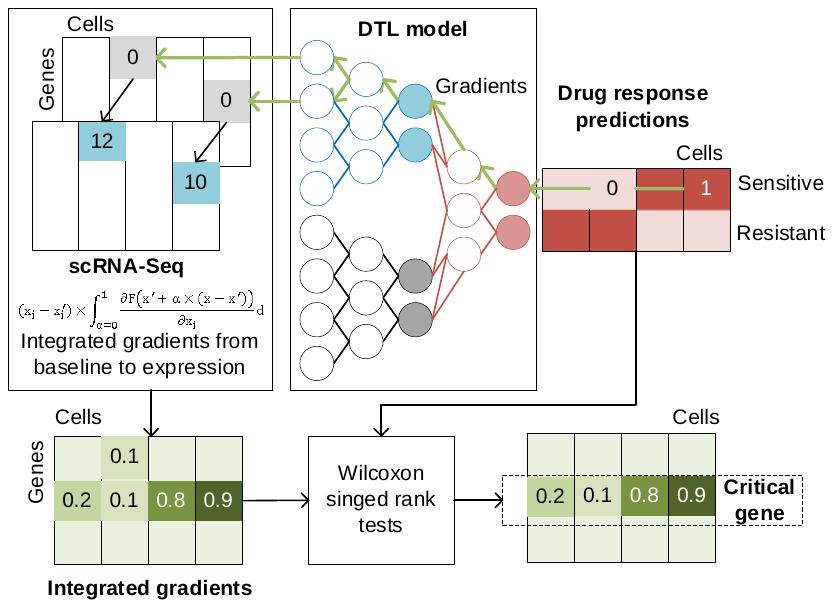

Supplementary Fig. S4. Procedure of the Integrated Gradient method on the gene interpretation from the DTL model.
